# IDENTIFICATION OF HUMAN PHOTORECEPTORS SUITABLE FOR CELL REPLACEMENT STUDIES IN A PRECLINICAL ACHROMATOPSIA MODEL

**DOI:** 10.64898/2026.06.22.733728

**Authors:** Patrick Schäfer, Andrea Corna, Thomas Kurth, Vincent Hain, Anne Schön, Susanne Ferguson, Andreea-Elena Cojocaru, Oriane Rabesandratana, Lucy Allan, Sarah Decembrini, Jesus Eduardo Rojo Arias, Olivier Goureau, Tiago Santos-Ferreira, Günther Zeck, Marius Ader

## Abstract

Cell replacement represents a potential treatment modality for retinal disorders characterized by photoreceptor loss. However, photoreceptor replacement approaches have not been clinically established. To take this forward, the main goal of this study was to systematically compare human photoreceptors of different ages and identify those that enable functional integration into the degenerative retina. Donor cells were isolated from iPSC-derived retinal organoids generated by a GMP-compliant protocol at differentiation days 120, 150, or 200 and transplanted subretinally into cone photoreceptor function loss 1 (Cpfl1) recipients, an inherited mouse model of cone degeneration. While younger photoreceptors showed slightly improved transplantation outcomes, donor photoreceptors of all culture stages displayed long-term survival, cone identity, structural integration into the host retina, and tight interactions with host Müller glia, including formation of a continuous outer limiting membrane. Transplanted photoreceptors showed signs of advanced maturation, including correct polarization with generation of apical inner- and outer segments, while basal synapses were formed with host bipolar cells. Electrophysiological assessment of host retinal ganglion cells revealed light-evoked responses in transplant-containing regions, providing evidence for functional incorporation of human photoreceptors into the mouse neuro-retinal circuitry. Thus, GMP-compliant human iPSC-derived photoreceptors are stable over a wide range of differentiation stages and constitute a robust cell source for retinal transplantation and functional repair. The findings provide important prerequisites for the development of standardized procedures towards clinical translation of photoreceptor replacement in the retina.

**Significant statement:** Cell transplantation represents a potential therapy for retinal dystrophies. However, conditions allowing stable functional integration and maturation of donor photoreceptors have not been defined. We systematically compared GMP-compliant iPSC-derived photoreceptors of different culture times after transplantation into a retinal degeneration mouse model. Photoreceptors of all ages showed structural integration and advanced maturation in close interaction with host Müller glia and bipolar cells, allowing restoration of light-driven responses. Thus human photoreceptors isolated from retinal organoids represent a robust source for clinical development of cell replacement in the diseased retina.

## INTRODUCTION

Loss of vision due to retinal disorders is a significant burden on quality of life ^1,2^. Age-related macular degeneration (AMD) and inherited retinal dystrophies (IRDs), such as retinitis pigmentosa or achromatopsia, are characterized by loss of light-sensing photoreceptors. Due to the ongoing aging of society and the high prevalence in the elderly, especially of AMD, the number of patients affected by such disorders is increasing rapidly ^3,4^. Therapeutic options are still limited, mainly advising changes of diet and lifestyle ^5–7^ or only slowing down disease progression ^8^. Despite enormous improvements in the development of several treatment strategies (e.g., gene therapy, small molecules, cell replacement, optogenetics, retinal prostheses), only one therapy, a gene therapy for an IRD (Leber’s congenital amaurosis (LCA) RPE65, Luxturna), is commercially available ^9^. Among the conceivable therapeutic options, replacement of lost photoreceptors is an attractive approach, allowing a single product to potentially restore lost anatomical structure and visual function independently of the disease etiology. Preclinical proof-of-concept studies confirmed the feasibility of photoreceptor transplantation into animal retinal degeneration models ^10^.

With the development of protocols for reprogramming human somatic cells into induced pluripotent stem cells (hiPSCs) ^11^, a new era in biomedicine started. Building on this pioneering work, a plethora of different protocols for human retinal organoid (HRO) generation have been developed within the last two decades ^12,13^. Today, hiPSCs are the preferred donor cell source over human embryonic stem cells (hESCs), as they are more easily accessible and ethically less restrictive. At the same time, multiple cell replacement approaches have been established and further developed utilizing PSC-derived photoreceptors, which are administered either as a single-cell suspension ^14–19^ or as sheets ^20–22^. This progress enabled the first human case study ^23^, and an increasing number of clinical trials are planned or already launched (NCT06789445/BlueRock Therapeutics; NCT06891885/Sumitomo Pharma America, Inc.; clinicaltrials.gov).

However, several hurdles remain on the way to clinical translation. First, the number of GMP-compliant protocols capable of generating high amounts of clinical-grade material is limited ^24,25,14^. Second, systematic assessment of optimal donor PR populations and side-by-side comparison of different HRO generation protocols are largely missing. Third, the proof of proper incorporation, including i) structural integration into the host retina, ii) correct polarization and maturation, iii) light-evoked function, and iv) long-term survival, is pending, incomplete, or not publicly available.

Here, we investigated the potential for functional integration of human iPSC-derived PR generated with a GMP-compliant protocol ^24^ after transplantation into a retinal degeneration mouse model. Transplantation of PR isolated at three different HRO ages (days in vitro (D)120, D150, D200) was compared following our standardized workflow ^16,26^. Key features of successful engraftment, including long-term survival, graft size, structural integration, interaction with host Müller glia, correct polarization, maturation, and graft-host connectivity, were assessed six months post-transplantation. Ultimately, we evaluated functional integration of donor photoreceptors by Multi-Electrode-Array (MEA) measurements.

## MATERIAL & METHODS

### Experimental animals

Adult *Cpfl1* mutant female mice (9–15 weeks of age) were used as recipients for cell transplantation ^27^. The *Cpfl1* mouse colony maintained in the CRTD animal facility was founded from mice provided by Bernd Wissinger (Institute of Ophthalmic Research, Tübingen, Germany). Mice were maintained on a 12-hour light/12-hour dark cycle with ad libitum access to food and water.

All animal experiments were approved by the ethics committee of the Dresden University of Technology and the Landesdirektion Dresden (approval numbers: TVV 38/2019). All relevant European Union regulations, German laws (Tierschutzgesetz), the Association for Research in Vision and Ophthalmology (ARVO) statement on the Use of Animals in Ophthalmic and Vision Research, and the NIH *Guide for the Care and Use of Laboratory Animals* (National Academies Press, 2011) were strictly followed for all animal work.

### hiPSC maintenance and differentiation of human retinal organoids

The previously characterized Crx-mCherry iPSC line ^15^ was used to differentiate retinal organoids. hiPSC were maintained in mTeSR^TM^ Plus (STEM CELL TECHNOLOGIES) with Rock Inhibitor and differentiated into retinal organoids using two distinct protocols previously described ^24,16^, see also supplementary methods.

### Human retinal organoid dissociation and photoreceptor isolation

HRO were dissociated with the Papain Kit (Worthington) according to Tessmer et al. 2023 ^26^. Briefly, retinal organoids were dissociated in 20 U/mL papain, followed by gentle titration with a fire-polished glass pipette and further processing according to the manufacturer’s instructions (Papain Dissociation System, Worthington). The cell pellet was resuspended in MACS buffer (0.5% BSA, 2 mM EDTA in PBS) to a concentration of approximately 5 million cells/mL. The cell suspension was filtered through a 35 μm mesh and kept on ice for FACS.

The single cell solutions were used to isolate PR with FACS (FACS AriaII, FACS AriaIII, FACS Aria Fusion, FACS Discover S8; BD), according to Tessmer et al. 2023 ^26^.

Briefly, the forward-scatter (FSC-A) and side-scatter (SSC-A) areas were used to discriminate cells from debris. Doublets were removed by gating the FSC height versus width and by the SSC height versus width. Dead cells were gated out using DAPI staining. Finally, mCherry^+^ cells were discriminated from auto-fluorescent cells using fluorescence detection at 505-525 nm (GFP) versus 579-594 nm (mCherry). After each FACSort, the isolated sample was re-analysed to assess and confirm purity and survival (not shown here).

### Subretinal transplantation

PR were transplanted based on Tessmer et al. 2023 ^26^. Following FAC sorting, purified cells were concentrated to 150,000 cells/µL, kept on ice and proceeded directly to transplantation. Note that transplantation occurred within a timeframe of 30 min - 3hrs post sorting to maintain cell viability. Mice were anesthetized via intraperitoneal injection of medetomidine hydrochloride (1 mg/kg, Domitor, Orion Pharma) and ketamine (30 mg/kg body weight; Ratiopharm) and pupils dilated with 2.5% phenylephrine/0.5% tropicamide (TU Dresden Pharmacy). The mouse head was secured and positioned with a head holder (Leica, Wetzlar, Germany) so that the optic nerve was visible in the central fundus. A hole was made in the ora serrata with a sharp 30-gauge needle (VWR, Germany). 1 µL of cell suspension then 0.2 µL of air was drawn into a 5 µl Hamilton syringe with a blunt 34-Gauge needle. The syringe was attached to a micromanipulator and transvitreally inserted into the eye under visual control aided by a microscope. The needle was placed nasally in the subretinal space, the air bubble quickly injected to create a bleb and then the cell suspension slowly injected. Directly following cell transplantation, 1 μL preservative-free triamcinolone acetonide suspension (80 μg/μL in NaCl prepared by the University Clinic Pharmacy, Dresden, Germany) was injected into the vitreous using a hand-held 10 μL Hamilton syringe with a blunt 34-gauge needle. Vitreal Triamcinolone injections were repeated on a monthly basis. Anesthesia was reversed by intraperitoneal injection of atipamezole (10mg/kg, Antisedan, Orion Pharma).

On each day of transplantation, referred to as transplantation round (N), one batch of HRO was used and transplanted into experimental animals (n) for one experimental paradigm D120, D150, and D200 (TP), and D200 (CTRL). All transplantation rounds were repeated resulting in the following total number of transplantation rounds (N) and transplanted eyes (n). D120 (TP): N = 2, n = 34; D150 (TP): N = 2, n = 31; D200 (TP): N = 2, n = 26; D200 (CTRL): N = 2, n = 20.

Out of those, representative samples were selected and further processed (detailed numbers are given in the figure legends).

### Tissue processing

Experimental animals were anesthetized with Isoflurane (Baxter) and sacrificed by cervical dislocation. Eyes were enucleated and fixed in 4% paraformaldehyde (PFA) solution for 1 hour at RT. The cornea and lens were then dissected away and the remaining eye cup cryopreserved in a graduated series of sucrose (10%, 30% and 50% (w/v) sucrose for 10 min, 30 min and 1 h respectively). Similarly, organoids were fixed in 4% PFA for 15 minutes at room temperature followed by cryopreservation. Eyes and organoids were embedded in NEG50 (Thermo Scientific), and cryo-sectioned to 12 µm thickness using the cryostat NX70 (Thermo Scientific), according to Gasparini et al. 2022 ^28^.

### Immunohistochemistry

IHC was performed based on Gasparini et al. 2022 with established markers ^28^. Briefly, frozen sections were air-dried briefly at room temperature (RT) and rehydrated for 15 minutes in phosphate buffered saline (PBS) followed by blocking (0.3% Triton X-100 solution with 1% bovine serum albumin (BSA) and 5% donkey serum) for 1 hour at RT. Sections were incubated overnight at 4°C with primary antibodies (Tab. 1) in blocking solution. Slides were washed with PBS three times to remove unbound primary antibody. Corresponding secondary antibodies (dilution of 1:1000) (Tab. 2), were incubated for 1 hour 30 minutes at RT together with 4ʹ,6-diamidino-2-phenylindole (DAPI; 1:15,000; Sigma).

**Table 1.** Primary antibodies.

| Antibody | Gene name | Cell type (location/structure) | Host | Company | Cat.no. | Dilution |
| --- | --- | --- | --- | --- | --- | --- |
| Arrestin 3 | ARR3 | cone (cytosol, OS) | gt | Novus Biologicals | NBP1-37003 | 1:100 |
| Glutamate-ammonia ligase | GLUL, GS | MG (cytosol) | ms | BD Biosciences | 610517 | 1:250 |
| Human mitochondria | - | human cells (mitochondria) | ms | Abcam | ab92824 | 1:500 |
| mCherry | - | n/a | ch | Merck Millipore | AB356481 | 1:1000 |
| Protein kinase C, alpha | PRKCA, PKCA | bipolar cells (cytosol) | rb | Santa Cruz Biotechnology | sc-208 | 1:300 |
| Recoverin | RCVRN | rods, cones, cone bipolar cells (cytosol, OS) | rb | Merck Millipore | AB5585 | 1:1500 |
| Solute carrier family 1 member 3 | SLC1A3, GLAST | MG, astrocyte (surface antigen) | ms | Miltenyi Biotec | 130-095-822 | 1:200 |
| Tight junction protein 1 | TJP1, ZO-1 | PR-MG and RPE (tight junctions) | rb | Thermo Fisher Scientific | 61-7300 | 1:200 |

legend: OS – outer segment, MG – Mueller glia, RPE – retinal pigment epithelial cells, gt – goat, ms – mouse, ch – chicken, rb – rabbit

**Table 2.** Secondary antibodies.

| Antibody | Reac-<br>tivity | Host | Company | Cat.no. | Dilution |
| --- | --- | --- | --- | --- | --- |
| Cy<sup>TM</sup>2 AffiniPure<sup>TM</sup> Donkey Anti-Mouse IgG (H+L) | ms | dk | Jackson<br>ImmunoResearch | 715-225-150 | 1:1000 |
| Cy<sup>TM</sup>3 AffiniPure<sup>®</sup> Donkey Anti-Chicken IgY (IgG) (H+L) | ch | dk | Jackson<br>ImmunoResearch | 703-165-155 | 1:1000 |
| Cy<sup>TM</sup>5 AffiniPure<sup>®</sup> Donkey Anti-Rabbit IgG (H+L) | rb | dk | Jackson<br>ImmunoResearch | 711-175-152 | 1:1000 |
| Cy<sup>TM</sup>2 AffiniPure<sup>TM</sup> Donkey Anti-Rabbit IgG (H+L) | rb | dk | Jackson<br>ImmunoResearch | 711-225-152 | 1:1000 |
| Cy<sup>TM</sup>5 AffiniPure<sup>®</sup> Donkey Anti-Goat IgG (H+L) | gt | dk | Jackson<br>ImmunoResearch | 705-175-147 | 1:1000 |
| Cy<sup>TM</sup>5 AffiniPure<sup>®</sup> Donkey Anti-Mouse IgG (H+L) | ms | dk | Jackson<br>ImmunoResearch | 715-175-150 | 1:1000 |
| Alexa Fluor<sup>®</sup> 488 AffiniPure<sup>®</sup> Donkey Anti-Goat IgG (H+L) | gt | dk | Jackson<br>ImmunoResearch | 705-545-147 | 1:1000 |

*legend: gt – goat, ms – mouse, ch – chicken, rb – rabbit, dk – donkey*

### Electron microscopy

For TEM or SEM eyes were collected from experimental animals as described for immunohistochemistry. After fixation in 4% FA in 100 mM phosphate buffer for 20-24 h, cornea, iris, lens and muscle tissue were cut away and the globe was stored in 1% FA/PB until further processing for EM as described ^29,28,30^.

### Ex vivo electrophysiological recordings of the mouse retina and related pharmacology

Recordings were performed using a 60-electrode glass microelectrode array (Multi-Channel Systems MCS GmbH) mounted beneath a microscope (BX 51, Olympus). Retinal dissections were performed according to previously described protocols ^31,32^. Dissections were performed under a stereomicroscope (MZ10 F Leica Microsystem) in dim red light. For transplanted eyes, the graft location was identified prior to dissection using fluorescence imaging under dim conditions. The corresponding retinal region was isolated and carefully positioned on the MEA, and graft alignment was verified via fluorescence microscopy.

For the experimental groups: D200 (TP) (TP HRO transplanted into *Cpfl1* mut), D200 (CTRL) (control HRO transplanted into *Cpfl1* mut, and non-tp (non-transplanted *Cpfl1* mut – age-matched control), the number of tested animals and samples included in the analysis were 9 animals/ 7 samples, 8 / 6, and 6 / 6, respectively. All procedures were approved by the Center for Biomedical Research, Medical University of Vienna, Austria.

Full field light stimulation was provided by a patterned OLED display (Sony ECX343E micro OLED display, Sony Semiconductor Solutions Corporation, Japan) focused via a 5× objective onto the retinal surface. Retinas were light-adapted for 30 minutes before each stimulus block at the respective background level. The individual light levels are provided in the supplementary method. After recordings, tissues were fixed in paraformaldehyde (PFA, Sigma) for immunohistochemistry.

Two pharmacological control experiments were conducted. During photopic stimulation: L-AP4 (50 µM; Tocris) was used to inhibit mGluR6-mediated transmission to ON bipolar cells, thereby also blocking rod-driven responses. ACET (100 µM; Tocris), applied in combination with L-AP4, silenced all photoreceptor synaptic output, serving as a negative control.

### Data acquisition and statistical analysis

#### Immunohistochemistry

The total graft area and inner segment (IS) formation was quantified using the ZEN 3.0 Blue Image Analysis Wizard of Axioscan Z-stack images. Therefore, every fourth serial retinal section throughout the whole eyecup of each sample was used. Obtained data was projected considering section thickness (12µm) and total number of series (4). The size (length) of each individual mCherry^+^ cell cluster was measured and analyzed for its level of incorporation. On the basis of the host ONL apical border, a line was drawn. The groups were defined as follows: isolated, 0% of the area below the apical border; partially integrated, 5%–80% below, fully integrated, greater than 80% below (Suppl. Fig. 2). Human inner segments protruded into the subretinal space (i.e., correctly formed) were counted manually based on human-specific hMito antibody (Tab. 1) and interrelated with size and integration status of the corresponding graft.

#### Electrophysiological recordings (MEA)

Recordings were acquired using the Multi-Channel Experimenter software (Multi Channel Systems MCS GmbH). Raw data were converted to .h5 format and preprocessed by applying a 4th-order Butterworth bandpass filter (300–4000 Hz), followed by subtraction of the median voltage to remove baseline offsets. Synchronization with the light stimulus was achieved via TTL triggers recorded on an analog Multi-Channel Experimenter input channel. Spike detection was performed using a median-based threshold-crossing method ^33^. Spiking activity was analyzed to quantify the percentage of responding ganglion cells, defined as electrodes exhibiting reproducible, stimulus-locked firing with a median firing rate across repetition, binned at 50 ms within 500 ms after light onset, exceeding 2.5 times the spontaneous rate measured before stimulation. Electrodes with fewer than one spike per minute were excluded, and retinas without light-evoked activity under any condition were removed from the analysis. The total percentage of responding cells was calculated by dividing the number of responding cells across all samples by the total number of detected cells across all samples. The Bias Index was computed for responding cells only and defined as BIAS INDEX = (number of spikes during light stimulation - number of spontaneous spikes) / (number of spikes during light stimulation + number of spontaneous spikes).

#### Image and data processing

Images and graphs were processed and generated using Zen 3.0 (Blue edition, Zeiss), Affinity Photo 2 & Designer 2 (Serif), and Prism 10 (GraphPad).

#### Statistics

Unless otherwise stated, data is displayed as mean +/- SEM and statistical significance is calculated using a one-way ANOVA with Tukey’s multiple-comparison test. A *P* value of less than 0.05 was considered as statistically significant.

## RESULTS

### Generation, transplantation and integration of human iPSC-derived photoreceptors into *Cpfl1* hosts

The integration potential of donor photoreceptors isolated from HROs of different ages were compared following transplantation into *Cpfl1* mice (Fig. 1A). HRO were generated from the photoreceptor-specific Crx-mCherry-iPSC reporter line ^15,16^ using a GMP-compliant protocol (Tenpoint (TP)) ^24^ and all experiments followed our standardized transplantation pipeline ^26^ including quality controlled (QC) HRO production, dissociation, FACS enrichment of mCherry^+^ PR and their subsequent subretinal transplantation (Fig. 1A). Given the specific TP protocol for HRO production that resulted in a distinct organoid-architecture including characteristic rosette formation (Suppl. Fig. 1A-C), control (CTRL) D200 HRO were additionally generated by an alternative protocol, that developed a laminated organoid structure (Suppl. Fig. 1D) ^29,30^. PR isolated from D200 (CTRL) HRO have been shown previously to extensively incorporate into the mouse retina after transplantation ^28^. Dissociation and subsequent enrichment of reporter-labelled photoreceptors (mCherry^+^) from all experimental groups, i.e., D120, D150, and D200 (TP), and D200 (CTRL) HRO, showed similar numbers of live cells per HRO (Fig. 1B), fraction of mCherry^+^ cells (Fig. 1C), viability after dissociation (Fig. 1D), viability after FACS isolation (Fig. 1E), and target cell yield (FACSorted/live cells for transplantation) (Fig. 1F).

**Figure 1:**
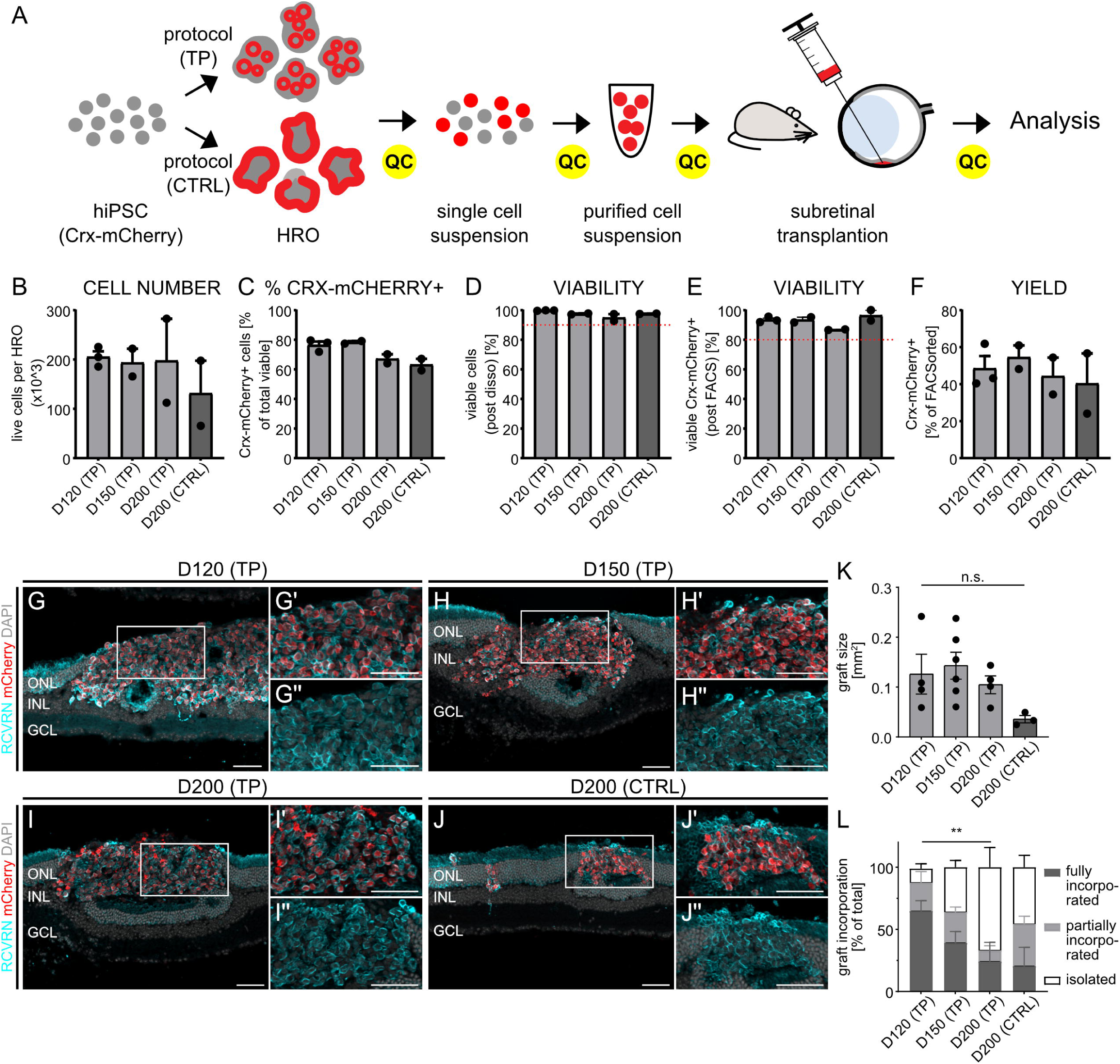
Transplanted photoreceptors of a wide range of age structurally integrate into the mouse retina and survive long-term. Experimental workflow: Crx-mCherry-hiPSC are differentiated into HRO using two protocols, Tenpoint (TP) vs control (CTRL), with defined age (TP: D120, D150, D200; CTRL: D200). HRO are dissociated, Crx-mCherry^+^ PR isolated by FACS and transplanted subretinally into a cone degeneration mouse model (Cpfl1 mouse). Six months (6M) post-transplantation eyes are collected and analyzed. This workflow is accompanied by a tight multi-step quality control (QC). This includes determination of live cell number per HRO (B), fraction of Crx-mCherry^+^ cells (C), viability after dissociation (D), viability after FACS isolation (E), and cell yield (FACSorted/live cells for transplantation) (F). B-F: N (HRO batches) per experimental paradigm: D120 (TP) N = 3, D150 (TP) N = 2, D200 (TP) N = 2, D200 (CTRL) N = 2 with 50 – 120 HRO used for each batch. Exemplary fluorescent images show capacity of donor cells isolated from Crx-mCherry reporter iPSC-retinal organoids at D120, D150, D200 (TP protocol), and D200 (CTRL protocol) to integrate into the retina of adult Cpfl1 hosts 6 months after transplantation (G-J). Human grafts (mCherry^+^, red) express the pan-PR marker RCVRN (cyan), and graft coverage (size) is not significantly different between the experimental paradigms (K). Donor PR mainly integrate into the ONL with transplants showing fully incorporated, partially incorporated or isolated phenotypes (L; see also Suppl. Fig. 3). Quantification is based on DAPI^+^ (nuclear dye)/mCherry^+^ transplant parts located within or outside the retina. The fraction of incorporated transplants decreases with age reaching significance when comparing D120 and D200 TP cells (L), indicating an age-dependent decline in capacity for integration. N (transplantation rounds), n (transplanted eyes) assessed per experimental paradigm: G) D120 (TP) N = 2, n = 12; H) D150 (TP) N = 2, n = 10; I) D200 (TP) N = 2, n = 6; J) D200 (Ader) N = 2, n = 11. For quantifications K-L) D120 (TP) N = 1, n = 4; D150 (TP) N = 2, n = 6; D200 (TP) N = 1, n = 4; D200 (CTRL) N = 1, n = 3. TP – Tenpoint, CTRL – control, mCherry – Crx-mCherry (pan PR reporter line), PR – photoreceptor, RCVRN – Recoverin (pan PR marker), ONL – outer nuclear layer, INL – inner nuclear layer, GCL – ganglion cell layer, statistics: one-way ANOVA (Tukey’s multiple comparisons), ** p-value < 0.01, scale bar 50µm

Following subretinal transplantation into the retinas of adult *Cpfl1* mice, human PR were identified by donor cell-specific expression of mCherry, and photoreceptor identity was confirmed by co-expression of the pan-photoreceptor marker recoverin (RCVRN) (Fig. 1G-J), with the vast majority of PR being cARR3^+^ cones (Suppl. Fig. 2A-D). For all experimental paradigms, i.e., donor cells isolated at D120, D150, and D200 (TP) or D200 (CTRL), PR were capable of integrating into host retinas and survived up to six months post-transplantation, demonstrating long-term survival (Fig. 1G-J). In all cases, integration of donor cells appears to occur by multicellular clusters rather than single cells. Moreover, while transplants predominantly occupied regions of the host outer nuclear layer (ONL), in rare cases donor photoreceptors also localized in the deeper inner nuclear layer (INL), but were not observed in the inner plexiform (IPL) or ganglion cell layer (GCL) (Fig. 1G-J). While no significant differences in graft size generated by the different donor photoreceptor ages were detected, a tendency for larger engraftment was observed in PR isolated from retinal organoids generated with the TP protocol in comparison to the CTRL D200 protocol (Fig. 1K). Integration level was assessed by measuring the amount of fully incorporated, partially incorporated, and isolated grafts (Suppl. Fig. 3), revealing a tendency for improved integration for the younger (D120, TP) photoreceptor fraction, which gained significance when compared to the D200 (TP) experimental group (Fig. 1L). Particularly, the proportion of fully incorporated transplants to isolated grafts, i.e., grafts that remain in the subretinal space without direct contact to the host retina (Suppl. Fig. 3), substantially increased with younger transplant age (Fig. 1L).

In summary, human photoreceptors isolated from three stages of retinal organoid culture and two different protocols show transplantation and integration potential in a mouse model of cone degeneration with a tendency of improved incorporation for younger photoreceptors. Particularly, the lamination of donor HROs (Suppl. Fig. 1A-D) appears not to be relevant for successful integration when using highly enriched PR suspensions for transplantation.

### Graft-host MG interaction

Close interaction of engrafted photoreceptors with host Müller glia (MG) appears to be highly important for their polarization and maturation ^16^. Therefore, immunohistochemical assessment of MG using a double antibody approach (GS/GLAST) ^28^ was performed (Fig. 2). Integrated grafts of all experimental paradigms showed close interaction with GS/GLAST^+^ processes, with many single donor photoreceptors appearing to be enwrapped by host MG (Fig. 2A-D and magnifications). Additionally, signs for a continuous outer limiting membrane (OLM) in areas with endogenous mouse- as well as human donor photoreceptors were observed (see intense fluorescent GS/GLAST^+^ line apically of the ONL), further providing evidence for structural integration of transplants into the recipient retinal tissue. In contrast, isolated grafts that remained in the subretinal space with no or limited contact to the host retina were not surrounded by MG processes.

**Figure 2:**
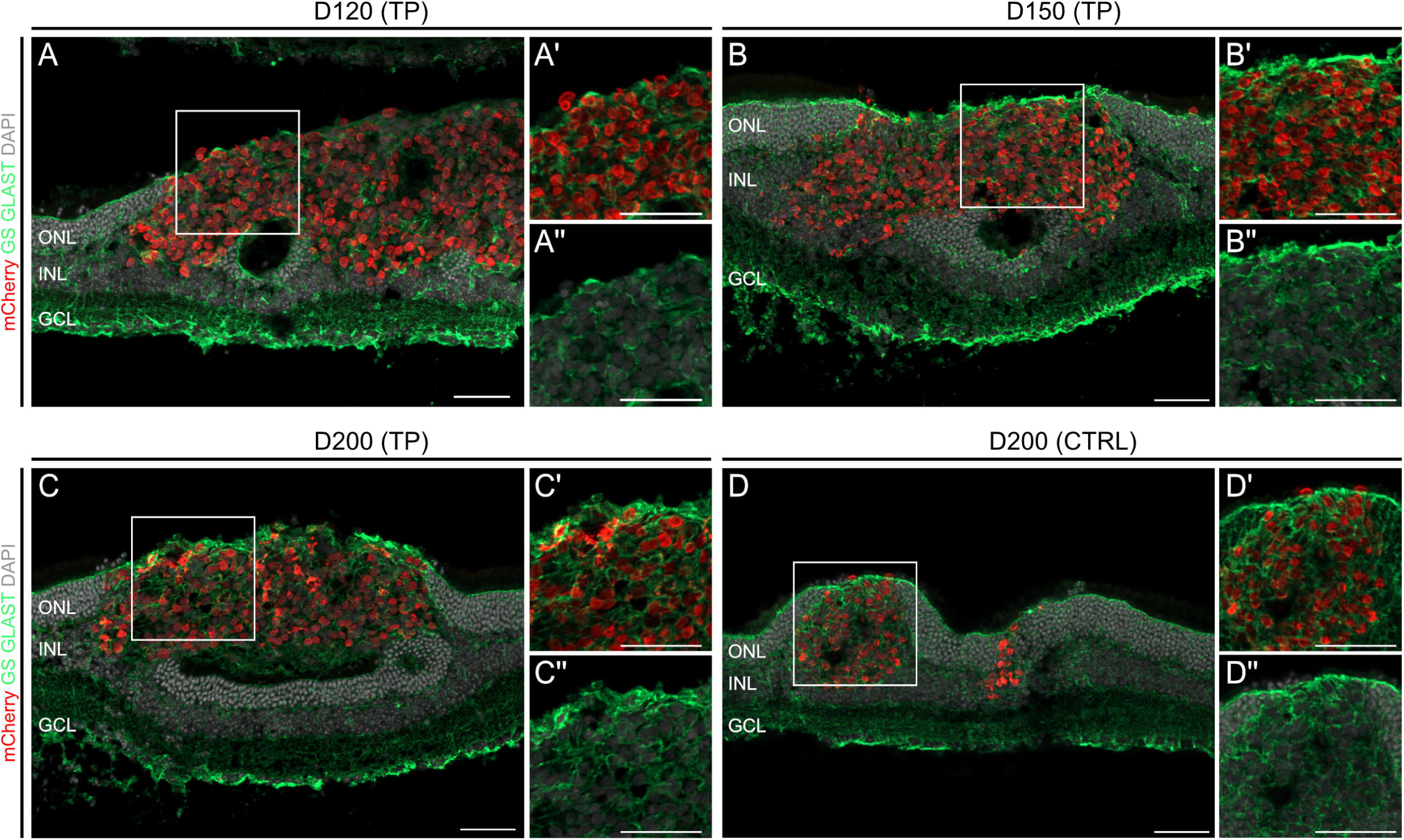
Donor photoreceptors tightly interact with host Müller glia cells. Exemplary fluorescent images (A-D) indicate close interaction between donor PR cells (mCherry^+^, red) and host MG (combination of the MG-specific markers GS and GLAST, green) in all experimental paradigms. Host MG processes (green) are tightly enwrapping donor cells (mCherry^+^). N (transplantation rounds), n (transplanted eyes) assessed per experimental paradigm. A) D120 (TP) N = 2, n = 9; B) D150 (TP) N = 1, n = 8; C) D200 (TP) N = 2, n = 4; D) D200 (CTRL) N = 2, n = 12. TP – Tenpoint, CTRL – control, PR – photoreceptor, MG – Müller glia cells, GS (GLUL) – glutamate-ammonia ligase, GLAST (SLC1A3)- solute carrier family 1 member 3, ONL – outer nuclear layer, INL – inner nuclear layer, GCL – ganglion cell layer, scale bar 50µm

### Photoreceptor polarization

Photoreceptor polarization, including formation of apical inner– (IS) and outer segments (OS) as well as basal synaptic terminals, depends on donor-host interplay and integration ^16^. IS formation was used as a read-out for polarization and maturation. Given the high concentration of mitochondria in IS, antibodies specific for human mitochondria (hMito) were used for immunohistochemical assessment of experimental retinas. Indeed, we identified mCherry^+^ PR of all experimental groups forming correctly polarized hMito^+^ protrusions, i.e., outgrowings into the subretinal space (SRS) towards the RPE (Fig. 3A-D, see magnified sections). Additionally, we sometimes also observed hMito^+^ protrusions that polarize into the lumen of inner graft rosettes. Interestingly, hMito^+^ protrusions were frequently localized apically of the OLM, visualized by the junction marker ZO1 (Fig. 3; also see Fig. 2 - OLM staining by GS/GLAST^+^), while mCherry^+^ nuclei/cell bodies remained basally of the OLM, typical for a polarized photoreceptor in the retina. Younger transplants (D120) showed a tendency for higher numbers of IS per mm^2^ graft than older transplants, though the quantified differences were not significant (Fig. 3E). In accordance to previous observations ^16^, in all experimental groups IS were particularly formed by transplants that were partially or fully incorporated (Fig. 3F, Suppl. Fig. 3), supporting the view that donor-host interaction is key for polarization and maturation of transplanted photoreceptors. In this line, highest propensity of IS formation (i.e., enhanced in vivo maturation) in D120 donor cells (Fig. 3E) correlated with the highest amount of graft integration (see Fig. 1L).

**Figure 3:**
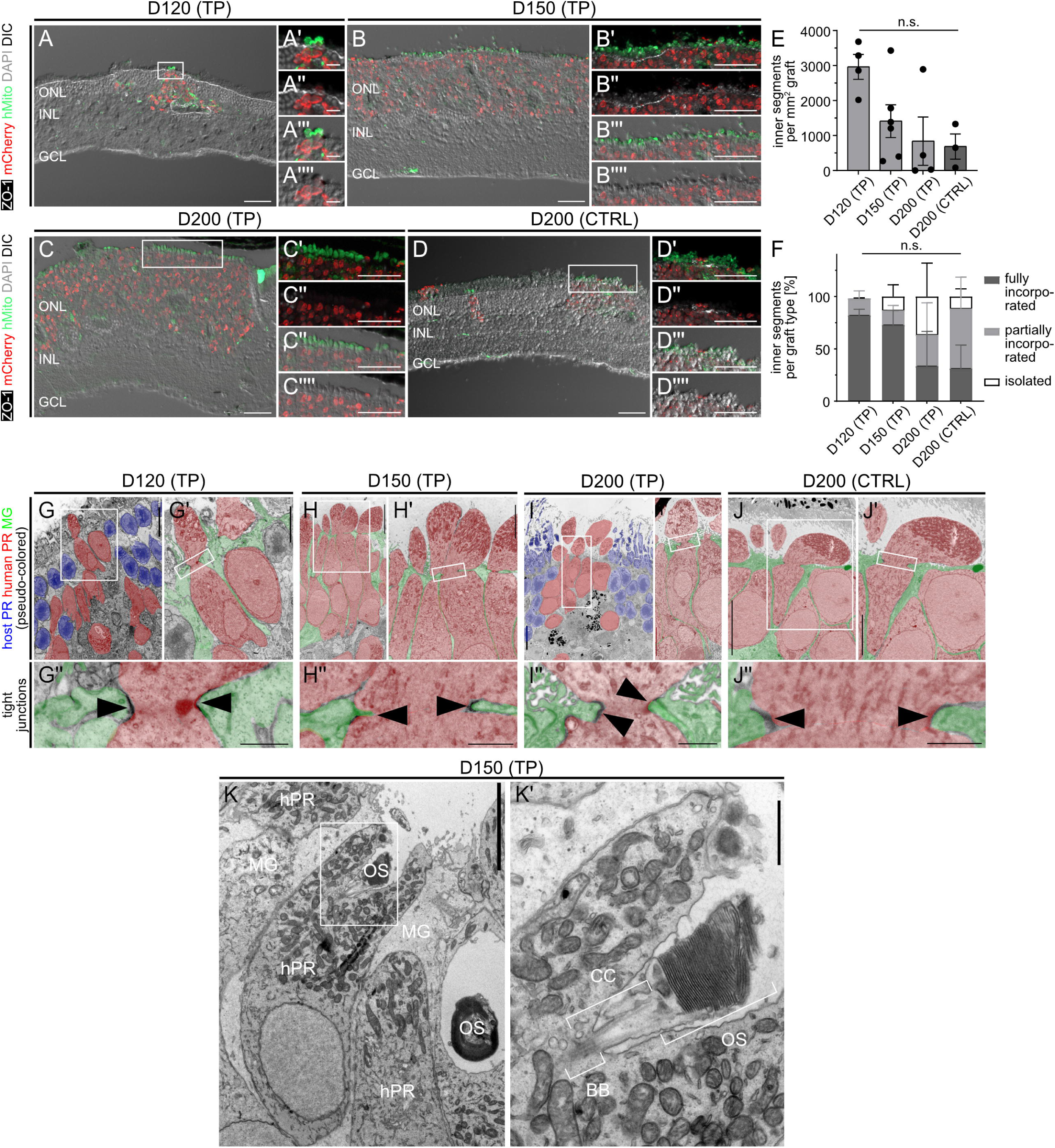
Donor photoreceptors polarize and form inner and outer segments. Exemplary fluorescent images (A-D) illustrate correct polarization of integrated donor PR with apical hMito^+^ IS-like protusions (green) and basal cell bodies (mCherry^+^) separated by a re-establishment OLM (ZO-1^+^, white) six months post transplantation. IS formation appears to decline with age of donor PR, without reaching significance (E). Apical hMito^+^ IS mainly form from integrated (fully or partially) grafts (F), indicating a need of donor-host interaction for PR polarization. TEM images verify incorporation into the host ONL and generation of apical, mitochondria-filled IS (G-J; donor PR pseudo-colored in red, host PR pseudo-colored in blue), with close interaction of donor PR with MG processes (G’, H-H’, I’, J-J’; MG pseudo-colored in green), including tight junctions formed at the base of donor IS (G’’-J’’, some labeled with arrow heads) indicative for OLM formation between donor cells and host MG. Example of an integrated donor PR with outer segment (OS, containing aligned membrane staples) apical of a mitochondria-filled IS, including connecting cilium (CC) and basal body (BB) (K, K’). N (transplantation rounds), n (transplanted eyes) assessed per experimental paradigm: A) D120 (TP) N = 2, n = 11; B) D150 (TP) N = 2, n = 9; C) D200 (TP) N = 2, n = 6; D) D200 (Ader) N = 2, n = 11. For quantifications E-F) D120 (TP) N = 1, n = 4; D150 (TP) N = 2, n = 6; D200 (TP) N = 1, n = 4; D200 (Ader) N = 1, n = 3. For EM, G) D120 (TP) N = 1, n = 3; H) D150 (TP) N = 2, n = 2; I) D200 (TP) N = 1, n = 1; J) D200 (CTRL) N = 1, n = 1. TP – Tenpoint, CTRL – control, hMito – human mitochondria, ZO-1 (TIP1) – zonula occludens-1/tight junction protein 1, IS – inner segment, PR – photoreceptor, OLM – outer limiting membrane, TEM – transmission electron microscopy, MG – Müller glia, PR – human photoreceptor, OS – outer segment, CC – connecting cilium, BB – basal body, ONL – outer nuclear layer, INL – inner nuclear layer, GCL – ganglion cell layer, statistics: one-way ANOVA (Tukey’s multiple comparison), scale bar (IHC): 50µm; scale bars (TEM): (G, H, I, J) 10µm, (G’-G’’, H’-H’’, I’-I’’, J’- J’’, K) 5µm, (K’) 1µm.

Ultrastructural analysis of experimental retinas by EM confirmed the formation of mitochondria-filled, apically polarized IS (Fig. 3G-J), as well as a close interaction with host MG, including formation of an OLM at the apical boundary (Fig. 3G-J). While mouse rod photoreceptors showed the typical inverted nuclear architecture (identified by the dark central area in the nucleus, Fig. 3G-J) ^34^, donor human cone nuclei were not inverted and appeared larger (Fig. 3G-J). In line, also IS of human transplants were larger with a different mitochondrial architecture than their mouse counterparts (Fig. 3G,I; also see Suppl. Fig. 4A-C for comparison of mouse and human PR). Furthermore, MG processes are found in close proximity to integrated transplants and form tight junctions to donor-derived IS (Fig. 3A-D, G-J). After six months in vivo, some donor PR further matured and also formed outer segments (OS) with characteristic structures including membrane staples, connecting cilium, and basal body (Fig. 3K; also see Suppl. Fig. 4D-G).

### Graft-host connectivity

Another key element for graft-mediated light detection is synaptic connectivity to host second-order interneurons, i.e., bipolar (BP) and horizontal cells. By using an ON BP marker (PKCa), neurite extensions of host BPs into transplants were observed in all experimental paradigms (Fig. 4A-D). Interestingly, EM analysis confirmed synapse formation by donor photoreceptors with characteristic features including synaptic ribbons and vesicles (Fig. 4E-H), as well as post-synaptic structures of second-order interneurons (Fig. 4E-H).

**Figure 4:**
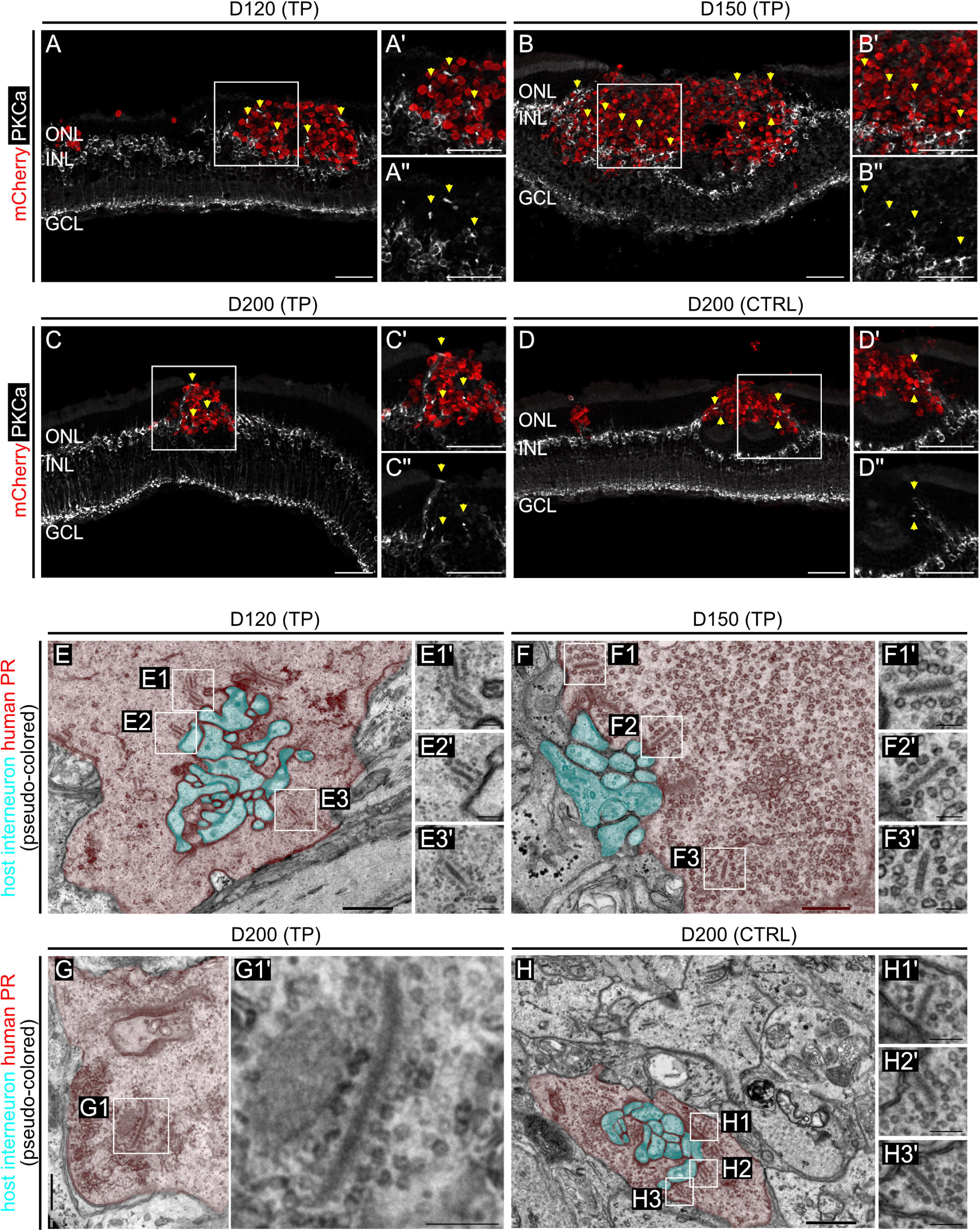
Donor cells integrate into the host neuronal circuit. Exemplary fluorescent images (A-D) indicate interaction between donor cells (mCherry^+^, red) and host secondary neurons, i.e., rod BP cells (PKCa^+^, white). Host BP extend neurites towards transplanted grafts and terminals (some marked by yellow arrowheads) are in close proximity to donor PR (A’-A’’, B’-B’’, C’-C’’). Selected TEM images confirmed synapse formation between donor PR (pseudo-colored in red) and host interneurons (pseudo-colored in cyan) (E, F, H). Higher magnifications show assembly of synaptic ribbons and accumulation of synaptic vesicles (E1’-E3’, F1’-F3’, G1’, H1’-H3’). N (transplantation rounds), n (transplanted eyes) assessed per experimental paradigm: A) D120 (TP) N = 2, n = 8; B) D150 (TP) N = 1, n = 8; C) D200 (TP) N = 2, n = 5; D) D200 (Ader) N = 2, n = 11. For EM, G) D120 (TP) N = 1, n = 3; H) D150 (TP) N = 2, n = 2; I) D200 (TP) N = 1, n = 1; J) D200 (CTRL) N = 1, n = 1. TP – Tenpoint, CTRL – control, BP – bipolar cells, PR – photoreceptors, TEM – transmission electron microscopy, PKCa (PRKCa) – protein kinase C alpha, ONL – outer nuclear layer, INL – inner nuclear layer, GCL – ganglion cell layer, scale bar (IHC): (A-D’’) 50µm; scale bar (TEM): (E, H) 1µm, (F, G) 500nm, (blowups, E1’-E3’, F1’-F3’, G1’, H1’-H3’) 200nm.

In summary, detailed histological and morphological analysis provided evidence for structural integration, maturation, and connections of transplanted human cone photoreceptors generated by a GMP-compliant protocol in cone-deficient mouse recipients, with signs of improved incorporation of D120 compared with older (D150, D200) transplants.

### Graft function

Functional integration of transplanted photoreceptors within the host retina was evaluated by Micro-Electrode-Array (MEA) measurements six months after transplantation (Fig. 5A). This approach enables direct measurement of retinal ganglion cell (RGC) activity in response to defined light stimulation protocols (Fig. 5B). The *Cpfl1* host model lacks functional cones but retains rod photoreceptors, which primarily mediate scotopic (low-light) vision. However, rods can remain responsive under photopic conditions at high light intensities ^35^. Our experiments were designed to identify conditions under which transplanted cones – as the main photoreceptor-type in transplants (Suppl. Fig. 2) - generate distinct responses compared with non-transplanted *Cpfl1* controls. Under photopic illumination, rod-mediated activation occurs only at high contrast and requires stronger stimuli to elicit responses than cone-mediated pathways ^35^. We therefore systematically varied stimulus contrast to test whether transplanted retinas exhibit enhanced responses at low contrasts, indicative of cone-mediated activity. As an initial metric of retinal responsiveness, we quantified the percentage of responsive cells, defined as the fraction of light-responsive RGCs, relative to the total number of detected cells. Both transplantation groups, D200 (TP) and D200 (CTRL), displayed a higher proportion of responsive cells at 33%, and D200 (TP) also at 44%, positive contrast compared to non-transplanted control retinas (Fig. 5C). These contrasts likely correspond to the range where rod responses are minimal while cone-driven activity is detectable, providing early evidence for functional integration of transplanted cones. At higher contrasts (≥ 66%), differences between transplanted and control retinas diminished (Fig. 5C), consistent with increased rod recruitment that masks cone-derived responses. Analysis of individual retinas at 33% and 44% stimulus contrast (Fig. 5D) further supported this observation: among the D200 (TP) samples (n = 7), four exhibited > 40% responsive cells, while three were comparable to controls; in the D200 (CTRL) group (n = 6), two retinas showed above-average responsiveness.

**Figure 5:**
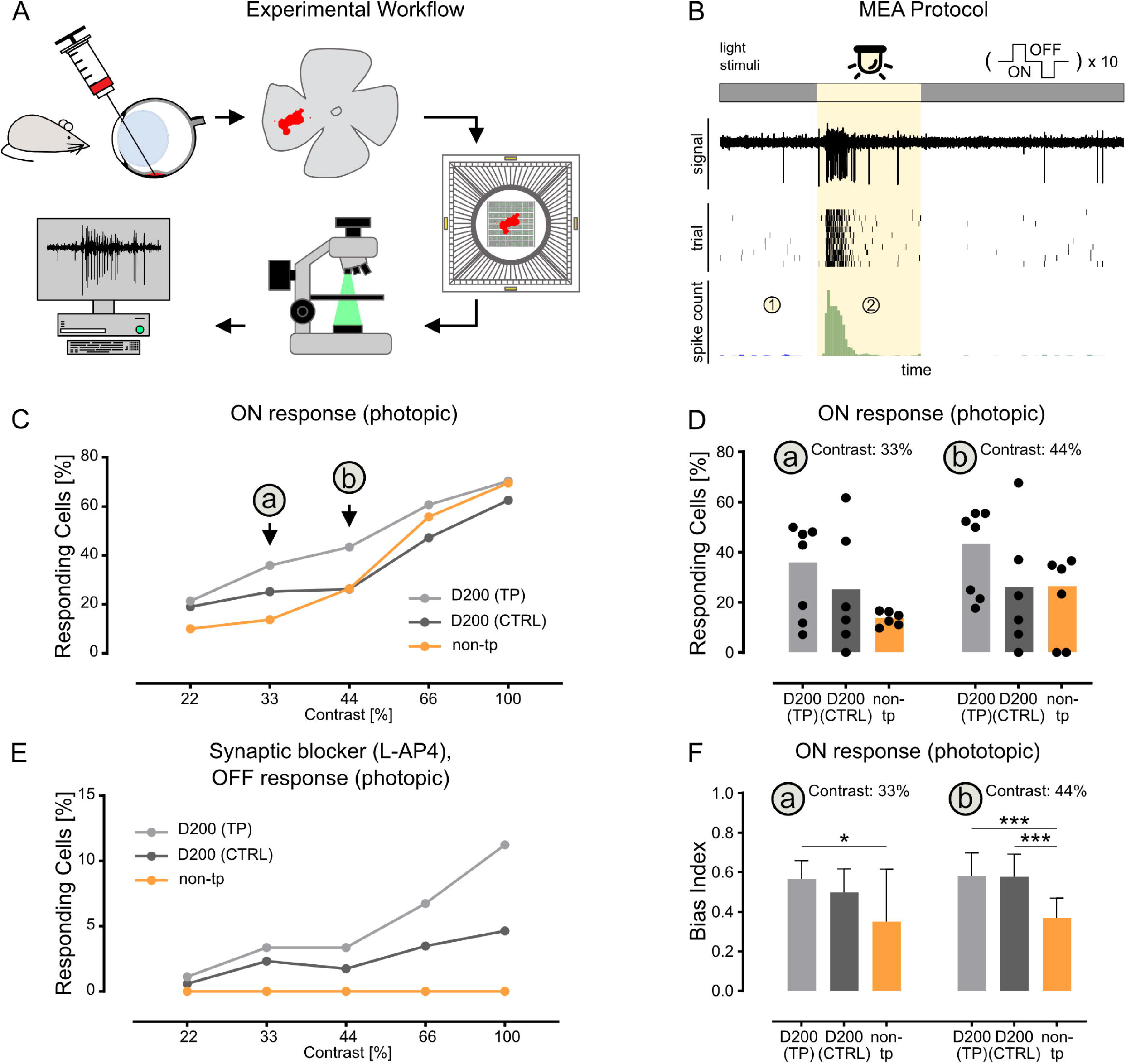
Functional integration of donor photoreceptors. Schematic of the experimental workflow for testing functional integration of transplanted photoreceptors using Micro-Electrode Arrays (MEAs) (A). A retinal sample containing the transplanted cell graft is isolated from the eye and placed on a MEA, where retinal responses to various light stimulation protocols are recorded from retinal ganglion cells (RGCs). Example of a light stimulation protocol (B). The retina is maintained under constant background illumination, and light stimuli are applied by increasing (ON stimuli) or decreasing (OFF stimuli) the light intensity at defined contrast levels. Each condition and contrast level is repeated 10 times. The MEA records RGC activity in the form of action potentials (spikes), which are classified as occurring before (1) or during (2) stimulation (B). Percentage of responding cells obtained by pooling all recorded retina samples for photopic (“daylight”) light stimulation at different contrast levels, ON response (C). Specifically at 33% (a) contrast, both transplantation groups, D200 (TP) and D200 (CTRL), show a higher percentage of responding cells compared to the control (non-transplanted) group. Percentage of responding cells at 33% and 44% contrast (as in panel C), shown for each retina individually (D). Percentage of responding cells at different OFF contrasts (i.e., decreases in light intensity relative to background) after application of the synaptic blocker L-AP4 (E). L-AP4 inhibits the ON response pathway in the retina, thereby isolating cone OFF responses from potential rod (ON-only) responses. Bias Index analysis for 33% and 44% photopic contrasts (F). The Bias Index quantifies the normalized increase in spike rate during illumination, (2) in panel B, relative to the baseline before light onset, (1) in panel B. A higher Bias Index indicates a stronger relative response to light stimulation. Data: C), E) responding RGC cells (pooled) as fraction of total. D) responding RGC (pooled) cells as fraction of total displayed as bars, data from individual samples displayed as dots, F) Bias index of individual responding RGC cells as median with 95% CI, statistic: Man-Whitney test, * p<0.05, *** p<0.001. N (transplantation rounds), n (transplanted eyes) used per experimental paradigm: C-F) D200 (TP) N = 2, n = 7; D200 (CTRL) N = 2, n = 6, non-transplanted (age-matched control) n = 6. TP – Tenpoint, CTRL – control, non-tp – non-transplanted (age-matched control), L-AP4 - L-2-amino-4-phosphonobutyric acid

To confirm the cone origin of the observed responses, we performed recordings in the presence of L-AP4, a selective group III mGluR agonist that blocks synaptic transmission between photoreceptors and ON bipolar cells, including rod bipolar cells ^16^. L-AP4 application abolished all ON light responses in control retinas and transplanted retinas (Suppl. Fig. 5A), confirming effective suppression of rod pathways. However, transplanted retinas retained measurable activity, with 2 to 11% (from ≥ 33% contrast) of cells remaining responsive to OFF stimuli, while responses to OFF stimuli in non-transplanted control retinas were fully depleted (Fig. 5E). This further confirmed the contribution and functional integration of transplanted human cones. Addition of L-AP4 and ACET (a kainate antagonist that blocks synaptic transmission between photoreceptors and OFF bipolar cells), which together abolish both rod-and cone-driven responses, completely eliminated light-evoked ON and OFF activity, thus confirming response specificity (Suppl. Fig. 5B,C).

Finally, response strength was assessed using the Bias Index by quantifying the normalized increase in spike rate during illumination relative to baseline activity (Fig. 5F). At low to intermediate contrasts (33 and 44%) - the same range where responsiveness was enhanced (Fig. 5C) - the Bias Index for ON stimuli was significantly higher in both D200 transplantation groups compared with controls (Fig. 5F). These results demonstrate that cone transplantation increases both the number of light-responsive RGCs and the magnitude of their responses, providing compelling evidence for functional integration of the transplanted cone photoreceptors into the host retina.

## DISCUSSION

Replacement of lost photoreceptors by transplantation of pluripotent stem cell (PSC)-derived donor photoreceptors represents a potential treatment option for regaining sight in patients suffering from retinal degenerative diseases. Recent preclinical studies provided evidence for functional integration of human photoreceptors into mouse models of retinal degeneration ^19,18,16,17,36^, laying the foundation for its further development. Indeed, a clinical phase 1 trial using photoreceptor suspensions has been started in 2025 (NCT06789445/BlueRock; clinicaltrials.gov) and photoreceptor sheet transplantation has been tested in two retinitis pigmentosa patients ^23^, with a clinical trial initiated (NCT06891885/Sumitomo Pharma America, Inc.; clinicaltrials.gov). However, published studies are highly heterogeneous in regard to used PSC lines, retinal organoid differentiation protocols, donor cell age, sorting technology, transplantation methodology, and showed high variety in donor photoreceptor integration, maturation, and functionality besides usage of different retinal degeneration mouse models ^17–19,28,36,37^. Thus, systematic comparisons might be of high importance to identify optimal conditions for successful photoreceptor transplantation. In this study, photoreceptors isolated from retinal organoids generated by a GMP-compliant protocol were isolated at different developmental stages for transplantation into a cone degeneration mouse model. While all donor cell fractions (D120, D150, D200) showed structural integration, proper maturation, and long-term survival for up to six months, younger donor cells (D120) appeared to have a more robust integration pattern by generating large transplants with improved maturation. Additionally, increased spiking responses of retinal ganglion cells underneath the transplants provided evidence for functional integration of human photoreceptors into the mouse neuro-retinal circuitry.

The retinal organoids generated using a GMP-compliant protocol showed major structural differences, particularly the formation of rosettes, in comparison to the laminated retinal layer in control organoids. Nevertheless, photoreceptors isolated from retinal organoids of both protocols and at different differentiation stages showed similar high viability and yield following dissociation and FACS enrichment, pointing to a general robustness of in vitro generated human photoreceptors independent of the used differentiation protocol. The chosen time points for PR isolation (D120, 150, 200) represent a wide range of retinal organoid development, but all stages were characterized by high numbers of Crx^+^ photoreceptors and low proliferation rates ^15,30^. Identification of suitable donor cells includes balancing cultivation time, with long enough time to generate optimal staged PR for integration after transplantation, and as short as possible culture times to reduce safety and cost issues. Indeed, no sign of uncontrolled growth or tumor formation was observed in any experimental eye. Our findings argue against a narrow time window for isolation of phenotypically stable donor PR from organoids for transplantation, spanning approximately 80 days (D120–200). However, it cannot be significantly expanded, as PR isolated from D250 retinal organoids showed significantly decreased integration potential ^16^. Whether earlier retinal organoid stages (<D120) can be used for the isolation of transplantable PR is currently unclear and has to be examined in future studies. Here, one has to take into account the significantly lower numbers of Crx^+^ PR and thus decreased yields after isolation, an increased amount of proliferating PR progenitor cells ^30^, as well as potential contamination with other retinal cell types. Indeed, recent studies providing protocols for accelerated generation of PR progenitors from PSCs within 24-32 days showed long-term survival after transplantation into animal models of retinal degeneration ^38,39^. However, only a subfraction of donor cells showed some expression of photoreceptor markers; structural integration appeared limited, but instead transplants formed tubulations and rosettes, and donor cells did not reflect the polarized morphology of mature photoreceptors, e.g., missing apical inner/outer segments and basal axonal terminals ^38,39^.

Interestingly, six months after transplantation, the vast majority of donor photoreceptors in our study were identified as cone photoreceptors, while the PR population of retinal organoids contained at least 50% rods/photoreceptor precursors at D120. Further studies are warranted to dissect the reasons for the low survival of engrafted rods following transplantation (process-related or host-related). Additionally, viability of rods vs. cones should be studied in further detail for the development of more targeted treatment approaches, e.g., rod-specific (retinitis pigmentosa) or cone-specific (AMD) retinal degenerative diseases. Moreover, in accordance with previous studies ^18,16,40^, no signs of material transfer between donor and host cells, i.e., no mCherry^+^ labeling of mouse cells, were observed. Regarding potential clinical applications and in line with a recent study ^41^, it is worth reporting that the used retinal organoids can be sent at around D100 alive via overnight courier in a temperature-controlled container and can be continued cultured for at least up to 100 days without losing their transplantation potential.

As previously shown ^16^, full donor cell integration supports graft polarization and maturation, which was also observed here. Donor cells without any contact to the host are not or rarely capable of polarizing and maturating, i.e., show proper IS formation. Indeed, for all used differentiation stages, structural integration with enwrapment by host Müller glia processes was observed. Accordingly, IS formation correlates with integration into the recipient tissue (see above); however, with a tendency of more IS to be generated in younger than older donor cells. It is not known yet whether this potential age-dependent decline is attributed to intrinsic differences (age-dependent capacity to cope with needs), to extrinsic cues during sample preparation (age-dependent changes in vulnerability and susceptibility to stress and damage) or to multiple etiologies. To elucidate this, the genomic signature of donor cells and its changes after transplantation need to be determined longitudinally. Importantly, IS formation was validated by ultra-structural analysis, confirming generation of polarized, mitochondria-filled IS, Müller glia processes around donor cell bodies, and tight junction formation between Müller glia endfeet and donor PR, underlining establishment of an OLM - an essential pre-requisite for PR polarization and barrier formation between ONL and inner-photoreceptor matrix.

Although PR polarization (apical IS and basal synapse formation) and signs for further maturation (formation of connecting cilia with attached OS-like structures) were observed, it appears not as complete as in vivo, given that the length of detected OS looks short and membrane staples not always fully aligned. Though transplanted PR maturate better within the mouse retina in vivo than remaining for the same time in retinal organoids in vitro ^16^, it simply might take even longer for complete maturation of transplanted PRs, though the total age of donor cells at the latest time point analyzed after transplantation was already more than one year (i.e., D200 + 6 months). This could be clarified in studies beyond six months after transplantation. Alternatively, the niche/tissue environment itself might limit further maturation. For example, the impact of mouse RPE on maturation of human PR and OS formation has not been systematically assessed up to now - maybe full human PR maturation requires interaction with human RPE, or even region-specific RPE subtypes ^42^. Also, the disease environment might influence maturation of donor photoreceptors, as interaction with Müller glia, including OLM and polarized IS formation, was rarely described in other studies using late-stage retina degeneration models ^20,18,43,44,36^.

An important step towards functional integration is synapse formation between donor photoreceptors and the remaining host circuit. Using orthogonal readouts, the formation of vesicle-containing ribbon synapses by donor photoreceptors was observed, including synaptic connections to host second-order neurons. Interestingly, bipolar cell dendrites appeared to grow into the transplants, forming ectopic synapses and did not form a clearly confined inner plexiform layer. While functional incorporation of donor PR was measured at the ganglion cell level by MEA, there is currently little knowledge of the extent to which such unconventional synapse formation influences signal transmission and processing ^10^. Thus, determining synapse formation and connectivity in more detail – i.e., how well they are rewired, in which quantities, also in regard to ON and OFF pathways – with correlation to functional data would be highly informative, also to fully interpret functional data.

Functional integration of human iPSC-derived cones into different host mouse retinas has been demonstrated by independent groups over the last few years ^16–18,36^. Functionality of human iPSC-derived cones within different host mouse retinas has been mainly demonstrated based on large-scale recordings from the *ex vivo* mouse retina using MEA ^16–19,36,45^, and in a few studies also by behavioral protocols ^19,18^. Indeed, our study demonstrates donor cone contribution to RGC ON and OFF responses, thus providing strong evidence of functional integration of human donor photoreceptors into the mouse retinal circuitry. While MEA recordings represent a highly sensitive read-out for functionality, transplantation studies so far used different light stimulation patterns for detecting distinct response types, i.e., some transplanted cells appear to integrate in the ON-, others in the OFF-retinal pathway, making comparisons of integration patterns between individual studies difficult. Thus, MEA protocols used so far may be carefully compared against each other to identify “standard operating procedures” to be followed in future experiments, allowing meta-studies regarding donor photoreceptor functionality under different conditions. This might be of particular interest to receive more insights into activation patterns that are indeed meaningful for image processing and vision restauration ^19^. Of course, this might be only the first step to be pursued, followed eventually by a more thorough classification into the functional retinal cell (sub)types known from retinal physiology ^46^.

### Conclusion

Human photoreceptors generated within PSC-derived retinal organoids appear to be a robust cell source for transplantation - usable over a large range of differentiation times, generated by different protocols, and surviving long-term after transplantation. Structural integration, including interaction with Müller glia and OLM formation, represent important prerequisites for proper polarization and IS/OS formation, while signal processing is conducted via synaptic connections to host bipolar cells and RGCs. Systematic assessment of donor cell age, pathologic environments, specified PR subtypes, and surgical methods should be conducted to improve donor PR survival, incorporation, and maturation to reveal optimized functional recovery. With the availability of diverse retinopathy mouse models and the upcoming of robust human disease models based on iPSC-derived retinal organoids ^29,47^, such comparison- and validation systems will be of utmost importance for the development of efficient cell replacement approaches for blinding diseases, which, after confirmation and adaptation in large animal models, will increase chances for successful translation towards clinical use.

## Acknowledgements

The authors like to thank members of the Flow Cytometry, Animal house, Electron-Microscopy, and Light Microscopy facilities at the CRTD and CMCB, Dresden University of Technology (TUD), for assistance with cell sorting, mouse maintenance, and imaging, respectively. Additional technical assistance was provided by Jochen Hentschel. The authors further thank Mai Thu Bui for her valuable assistance with the experimental procedures. They also acknowledge the Core Facility for Laboratory Animal Breeding and Husbandry (CFL) at the Medical University of Vienna for their support.

## Data Availability Statement

This study did not generate new unique reagents or new code. Requests for further information and resources should be directed to and will be fulfilled by the lead contact, Marius Ader.

## Author contributions

**Patrick Schäfer** (Conception and design, administrative support, collection and assembly of data, data analysis and interpretation, manuscript writing, final approval of manuscript)

**Andrea Corna** (Conception and design, collection and assembly of data, data analysis and interpretation, manuscript writing)

**Thomas Kurth** (collection and assembly of data, data analysis and interpretation)

**Vincent Hain** (collection of data)

**Anne Schön** (Provision of study material)

**Susanne Ferguson** (Provision of study material)

**Andreea-Elena Cojocaru** (collection and assembly of data)

**Oriane Rabesandratana** (Provision of study material)

**Lucy Allan** (Provision of study material)

**Sarah Decembrini** (Provision of study material)

**Jesus Eduardo Rojo Arias** (Provision of study material)

**Olivier Goureau** (Provision of study material, final approval of manuscript)

**Tiago Santos-Ferreira** (Conception and design, financial support, administrative support, final approval of manuscript)

**Günther Zeck** (Conception and design, manuscript writing, final approval of manuscript)

**Marius Ader** (Conception and design, financial support, administrative support, manuscript writing, final approval of manuscript)

## Supplemental Information

Document S1. Figures S1–S5

