## Supplemental Info for "IDENTIFICATION OF HUMAN PHOTORECEPTORS SUITABLE FOR CELL REPLACEMENT STUDIES IN A PRECLINICAL ACHROMATOPSIA MODEL"

### SUPPLEMENTAL MATERIAL

### Supplementary methods

#### Differentiation of retinal organoids – protocol 1 (Tenpoint Therapeutics)

Briefly, undifferentiated hiPSC were plated on vitronectin coated dishes and culture with mTeSR™ Plus with ROCK inhibitor (see table 3) until they reached 80% confluence (day 0 of differentiation). On the same day (day 0), cell culture medium was replaced for CTS™ Essential 6 medium for 24 hours and replaced by Neural Induction Medium (NIM, see table 4). 50% of the NIM medium was replaced every second day from day 2 to day 28 of differentiation. On day 28, a G23 needle attached to a syringe was used to pick optic vesicle-like structures (OVs). Using the needle's tip a square frame was outlined as close as possible to every OV without disturbing it and gently removed the detaching OVs. OVs were then cultured in floating conditions in Retinal Maturation Medium (RMM, see table 5) supplemented with 10ng/mL human recombinant FGF2 (R&D Systems; 234-GMP-025) in a 6-well low binding plate until day 35. Human retinal organoids (HROs) were further cultured in RMM until the day of transplantation.

**Table 3: General reagents for hiPS cell maintenance and dissociation**

| Product | Supplier | Reference |
|---|---|---|
| GCDR | Stem Cell Technologies | 100-0485 |
| mTeSR™ Plus, GMP | Stem Cell Technologies | 100-0274 and 100-0275 |
| Rock Inhibitor Y-27632 | TOCRIS | TB1254-GMP |

**Table 4: Composition of Neural Induction Medium (NIM)**

| Product | Supplier | Reference |
|---|---|---|
| CTS Essential 6 Medium | Thermo Fisher Scientific | A4238501 |
| (100X) CTS™ (Cell Therapy Systems) N-2 Supplement | Thermo Fisher Scientific | A1370701 |

**Table 5: Composition of Retinal Maturation Medium (RMM)**

| Product | Supplier | Reference |
| --- | --- | --- |
| Gibco™ CTS™ KnockOut™ DMEM/F-12 | Thermo Fisher Scientific | A1370801 |
| CTS™ GlutaMAX™-I supplement | Thermo Fisher Scientific | A1286001 |
| 1% MEM non-essential amino acids | Thermo Fisher Scientific | 11140050 |
| CTS™ B-27™ Supplement XenoFree (50X) | Stem Cell Technologies | A5047501 |
| 10 units/ml Penicillin + 10 *μ*g/ml Streptomycin | Thermo Fisher Scientific | 15140122 |
| 10 ng/ml of recombinant human FGF2 | R&D | 233-GMP-025 |

#### Differentiation of retinal organoids – protocol 2 (Ader lab)

The Crx-mCherry iPSC line [47] was maintained in mTeSR1 (STEMCELL Technologies) on Matrigel-coated plates and split using ReleSR at room temperature (STEMCELL Technologies). Stem cells were differentiated into retinal organoids using a previously described optimized protocol [29].

Briefly, when hiPSC reached 80% confluence (Crx-mCherry) these were detached to small cell clusters with ReLeSR and resuspended in Matrigel (growth-factor reduced, BD Biosciences). Matrigel was allowed to gel for 5-7min at room temperature and was then gently dispersed into small clumps in N2B27 media (1:1 DMEM/F12: Neurobasal A media, 1% B27+VitaminA, 0.5% N2, 1% penicillin/streptomycin, 1% GlutaMAX, 0.1 mM 2-mercaptoethanol, see table 6). This suspension was added to 6-well ultra-low attachment plates (Nunclon Sphera, Thermo Fisher) allowing floating neuroepithelial cysts to form within the first few days of culture. Cysts were then plated on Matrigel coated plates on day 5, followed by gentle Dispase detachment (Stem cell technologies) on day 13. Detached clusters were transferred to 9 cm ultra-low attachment plates (Nunclon Sphera, Thermo Fisher) in B27 media (DMEM/F12, 1% B27 without Vitamin A, 1% penicillin/streptomycin, 1% GlutaMAX, 1% NEAA, 0.1% Amphotericin B, see table 7). On day 30, retinal epithelial domains were manually isolated using surgical tweezers (Fine Science Tools, Dumont No. 5). From D25, B27 media was supplemented with 10% FBS (see table 8) and from D100 N2+FBS media was used (DMEM/F12, 1% N2, 10% FBS, 1% penicillin/streptomycin, 1% GlutaMAX, 0.1% Amphotericin B, see table 9). Synthetic retinoid analogue EC23 (0.3 μM) was supplemented from D25 to D120. Media was refreshed with 50% media change every 2-3 days from day 13 onwards.

**Table 6 Composition of N2B27 medium (Ader protocol) (D0-12)**

| Product | Supplier | Reference |
|---|---|---|
| DMEM/F-12, GlutaMAX™ Supplement | Thermo Fisher Scientific | 31331-028 |
| Neurobasal™ Medium | Thermo Fisher Scientific | 21103-049 |
| B-27™ Supplement (50x) (w/ vit.A) | Thermo Fisher Scientific | 17504-044 |
| N-2 Supplement (100x) | Thermo Fisher Scientific | 17502-048 |
| Pen/Strep | Sigma-Aldrich | P0781 |
| GlutaMAX™ Supplement | Thermo Fisher Scientific | 35050061 |
| 2-Mercaptoethanol (50 mM) | Thermo Fisher Scientific | 31350010 |

**Table 7 Composition of B27 medium (Ader protocol) (D13-24)**

| Product | Supplier | Reference |
|---|---|---|
| DMEM/F-12, GlutaMAX™ Supplement | Thermo Fisher Scientific | 31331-028 |
| B-27™ Supplement (50x) (w/o vit.A) | Thermo Fisher Scientific | 12-587-010 |
| Pen/Strep | Sigma-Aldrich | P0781 |
| NEAA | Thermo Fisher Scientific | 11140035 |
| Amphotericin B (Fungizone) | Thermo Fisher Scientific | 15290026 |

**Table 8 Composition of B27 medium + FBS (Ader protocol) (D25-99)**

| Product | Supplier | Reference |
|---|---|---|
| DMEM/F-12, GlutaMAX™ Supplement | Thermo Fisher Scientific | 31331-028 |
| B-27™ Supplement (50x) (w/o vit.A) | Thermo Fisher Scientific | 12-587-010 |
| Pen/Strep | Sigma-Aldrich | P0781 |
| FBS | Thermo Fisher Scientific | A5256801 |
| NEAA | Thermo Fisher Scientific | 11140035 |
| Amphotericin B (Fungizone) | Thermo Fisher Scientific | 15290026 |
| EC23 | Tocris | 4011 |

**Table 9 Composition of Retinal Maturation Medium (RM2) (Ader protocol) (D100-200)**

| Product | Supplier | Reference |
|---|---|---|
| DMEM/F-12, GlutaMAX™ Supplement | Thermo Fisher Scientific | 31331-028 |
| N-2 Supplement (100x) | Thermo Fisher Scientific | 17502-048 |
| Pen/Strep | Sigma-Aldrich | P0781 |
| FBS | Thermo Fisher Scientific | A5256801 |
| Amphotericin B (Fungizone) | Thermo Fisher Scientific | 15290026 |
| EC23 (until D120) | Tocris | 4011 |

#### Quality and control (QC)

Each HRO differentiation protocol is capable of generating high numbers of photoreceptors (Fig. 1C, Suppl. Fig.1A-D). The percentage of Crx-mCherry$^+$ cells is in a similar range (58-80%) (Fig. 1C).

*Our workflow is very robust and lead to high quality of donor cells. Each stage of sample generation and preparation is equipped with a QC measure that allows exclusion of donor cells not appropriate for the laborious downstream procedures (*Fig. 1A*). All samples passed our internal quality criteria; viability after HRO dissociation (*Fig. 1*D, red dashed line), after FACSort (*Fig. 1E, red dashed line*), and cell yield/recovery (*Fig. 1F*). As QC measure for the HRO culture, we monitor cell number per HRO (Fig. 1B) and the fraction of Crx-mCherry*$^+$ *cells (Fig. 1C). So, transplantations fail solely due to uncontrollable events or the biology itself. By integrating a tight QC we reduced variability of transplantations.*

#### Transmission Electron Microscopy (TEM)

For TEM, the samples were further dissected to small pieces (0.5-1 mm) with the transplanted cells under a fluorescent stereo microscope (Leica MZ10F, Leica Microsystems, Wetzlar, Germany). Tissue pieces were postfixed at least overnight in modified Karnovsky's Fixative (2% glutaraldehyde, 2% paraformaldehyde in 100 mM phosphate buffer) at 4°C. Samples were further postfixed and contrasted following the OTO-procedure by incubating them in a 2% $OsO_4$ solution (2% OsO4, 1.5% potassium ferrocyanide, 2 mM CaCl2 in water), then in 1% aqueous thiocarbohydrazide, followed by another osmium step in 2% aqueous $OsO_4$ with washes in water between all incubation steps [48]. Finally, the samples were contrasted with 1% uranyl acetate in water, and washed several times in water. Contrasted samples were dehydrated in a graded ethanol series from 30% to 100% on a molecular sieve (30%, 50%, 20 min each, 70%, 90%, 95%, 3x 100%, 30 min each), before embedding in epoxy resin EMBed 812/Araldit 1:1) through stepwise infiltration (25%, 50%, 75% resin in water-free ethanol for 1h each, pure resin overnight, pure resin for 5 hrs) and curing at 65°C overnight.

To find the graft region, semithin sections were prepared with a Leica UC6 ultramicrotome (using glass knifes) and stained with toluidine blue/borax. Semithin sections were imaged with a Keyence Biozero BZ8000. For TEM, 70 nm ultrathin sections were cut with the ultramicrotome (using a diamond knife from Diatome), collected on formvar-coated slot grids and contrasted once more with lead citrate [49] and uranyl acetate. Samples were then imaged with a Jeol JEM1400 Plus transmission electron microscope (JEOL, Freising, Germany; camera: Ruby, JEOL) running at 80 kV acceleration voltage.

#### Scanning Electron Microscopy (SEM)

For SEM, samples were further dissected and the RPE/choroid/sclera layers were separated from the retina to generate en face views on the retina and/or RPE surfaces. The dissected samples were postfixed in modified Karnovsky (2% glutaraldehyde and 2% formaldehyde in 100 mM phosphate buffer), washed 2× 5 min with PBS and 3 × 5 min with bi-distilled water, poststained with 1% osmium tetroxide ($OsO_4$) for 2 hrs on ice, washed several times in water, postfixed in 1% aqueous uranyl acetate and dehydrated in a graded series of ethanol/water mixtures up to pure ethanol (30%, 50%, 70%, 96% and 3 × 100% on molecular sieve, 15min each). The samples were dried via critical point drying with a Leica CPD300 (Leica Microsystems, Wetzlar, Germany). Dried samples were mounted on 12 mm aluminium stubs using conductive carbon tabs and additionally grounded with conductive liquid silver paint. To increase contrast and conductivity, samples were sputter coated with gold using the BAL-TEC SCD 050 sputter coater (settings: 80 sec, with 60 mA, at 5 cm working distance). Finally, samples were imaged with a JSM 7500F scanning electron microscope (JEOL, Freising, Germany) running at 5kV (lower SE-detector, working distances between 7.5 and 8 mm).

### Ex vivo electrophysiological recordings of the mouse retina and pharmacology

#### Micro-Electrode Array

Recordings were performed using a 60-electrode glass microelectrode array (MEA; 30 µm diameter, 200 µm spacing, 3 × 3 mm area; 60MEA200/30iR-ITO, Multi-Channel Systems MCS GmbH) mounted on a MEA60-System headstage beneath a microscope (BX, Olympus). Each MEA was first cleaned with a detergent solution (5% Tickopur R60Stamm/Berlin), plasma treated (ZEPTO, Diener electronic), and coated with poly-L-lysine (200 µL, 1 mg/mL, P1399, MW 150–300 kDa, Sigma-Aldrich) to improve retinal adhesion.

#### Retina preparation

Retinal dissections were performed according to previously described protocols [31,32], with adaptations for identifying the graft region in transplanted samples. Recordings were performed on transplanted *Cpfl1* mut mice (see the transplantation section for details). In addition, retinas from non-transplanted control animals (*Cpfl1* mut) were also included. Briefly, retinal preparation involved removal of the cornea and lens, followed by hemisecting the eye to isolate the retina. The vitreous was carefully removed, and a ~3–4 mm² retinal segment was placed ganglion cell side down onto the MEA. All samples were maintained in carbogenated Ames' medium (95% $O_2$, 5% $CO_2$; MilliporeSigma), supplemented with $NaHCO_3$. Dissections were performed under a stereomicroscope (MZ10 F Leica Microsystem) in dim red light. For transplanted eyes, the graft location was identified prior to dissection using fluorescence imaging under dim conditions. The corresponding retinal region was isolated and carefully positioned on the MEA, and graft alignment was verified via fluorescence microscopy. Transplanted retinas that did not display any detectable fluorescent graft signal were not recorded. For the experimental groups: D200(TP) (TP HRO transplanted into *Cpfl1* mut), D200 (CTRL) (control HRO generated with Ader protocol, transplanted into *Cpfl1* mut), and non-tp (non-transplanted *Cpfl1* mut – age-matched control), the number of animals tested and samples included in the analysis were as follows: 9 animals / 6 samples, 8 / 6, and 6 / 6, respectively.

All animal procedures were approved by the Center for Biomedical Research, Medical University of Vienna, Austria.

#### Light stimulation

Full field light stimulation was provided by a patterned OLED display coupled via a 5× objective onto the retinal surface. Retinas were light-adapted for 30 minutes before each stimulus block at the respective background level: scotopic 0.02629 µW/mm², followed by photopic at 0.46749 µW/mm². For each background (scotopic and photopic), we tested 5 light contrast levels (22, 33, 44, 66, 100% Michelson contrast). Each one of the contrasts included (i) 10 repetitions of: bright contrast and dark contrast (2 s) intervalled by a break (background light intensity for 4 s) (see below). After recordings, tissues were fixed in paraformaldehyde (PFA, Sigma) for immunohistochemistry.

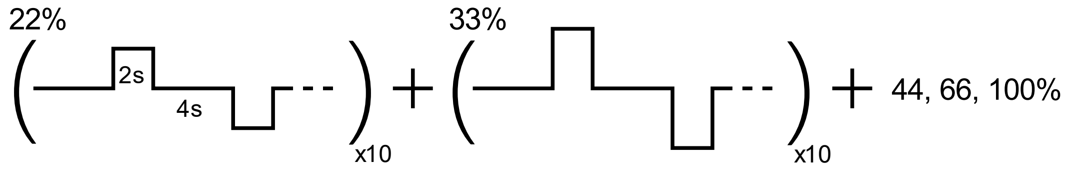


Light stimulation protocol for a defined background light condition.

#### Pharmacology

In addition to the scotopic and photopic stimulation protocol, two pharmacological control experiments were conducted following the same protocol of the photopic regime with the addition of the blocker at the beginning of the adaptation phase. To isolate specific retinal responses, two pharmacological blockers were applied during photopic stimulation: L-AP4 (50 µM, Tocris) was used to inhibit mGluR6-mediated transmission to ON bipolar cells, thereby also blocking rod-driven responses. ACET (100 µM, Tocris), applied in combination with L-AP4, silenced all photoreceptor synaptic output, serving as a negative control.

#### Supplemental figures

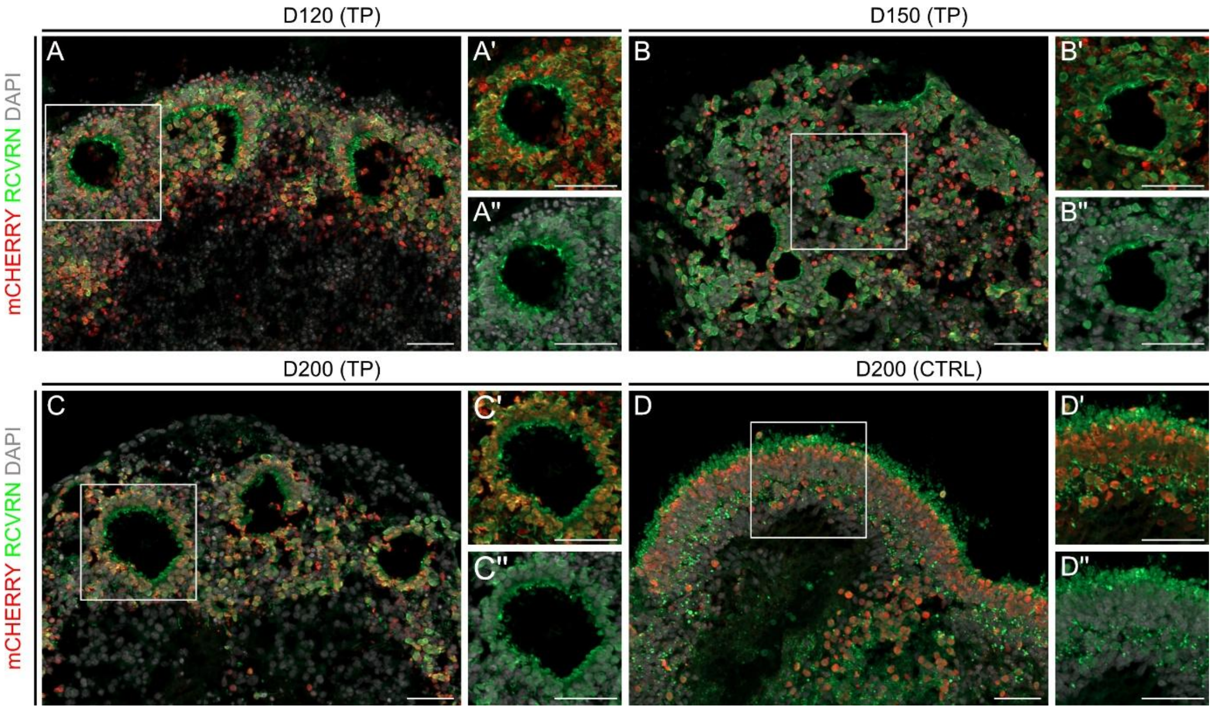


***Suppl. Figure 1: Structure of retinal organoids generated by TP or CTRL protocols***

*Exemplary fluorescent images of sectioned Crx-mCherry-iPSC-derived HRO (A-D). All organoids generate high numbers of mCherry$^{+}$ (red) PR co-expressing the pan-PR marker recoverin (RCVRN, green). Tenpoint (TP) HROs at D120, D150, or D200 are characterized by rosette formation (A-C), the region where the majority of PR are located (A'-A'', B'-B'', C'-C''). In contrast, CTRL HRO at D200 show a laminated structure including an ONL-like layer mainly populated by PR (D) - a structure highly similar to the in vivo retina.*

*hiPSC – human induced pluripotent stem cells, HRO – human retinal organoid, D – days in culture, TP – Tenpoint, CTRL – control, RCVRN – Recoverin, scale bar: (A-D'') 50µm*

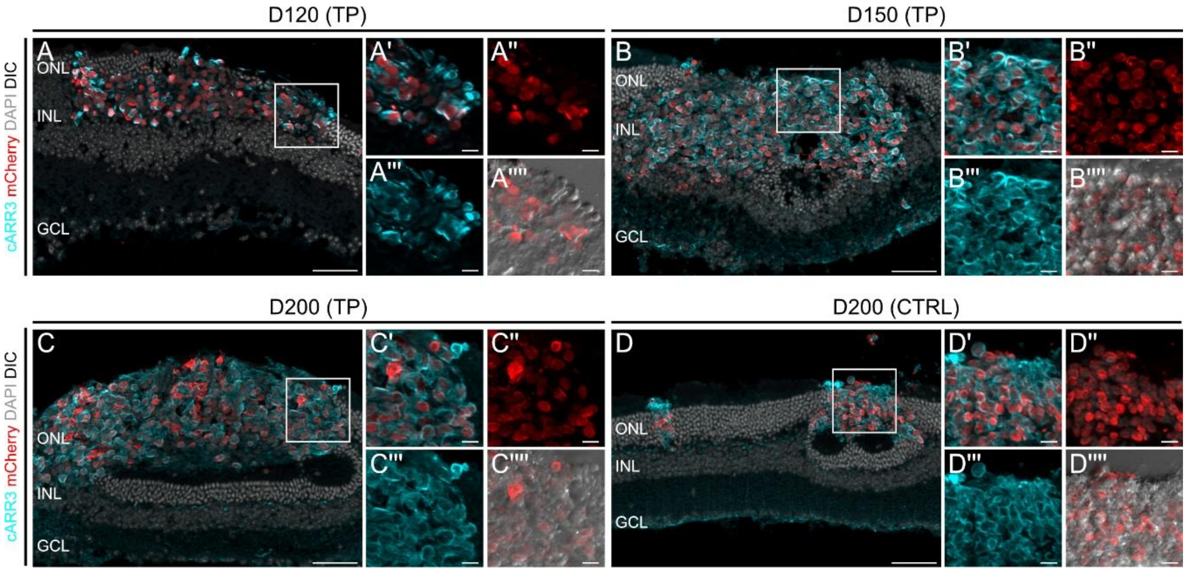


***Suppl. Figure 2: Six-months post transplantation grafts are enriched in cone photoreceptors***

*Exemplary fluorescent images of all experimental paradigms demonstrate that the vast majority of Crx-mCherry$^{+}$ (red) grafted cells 6M post-transplantation express the cone-specific marker cARR3$^{+}$ (cyan). N (transplantation rounds), n (transplanted eyes) assessed per experimental paradigm: A) D120 (TP) N=2, n=9; B) D150 (TP) N=1, n=8; C) D200 (TP) N=2, n=5; D) D200 (CTRL) N=2, n=11.*

*TP – Tenpoint, CTRL – control, mCherry – Crx-mCherry (pan-PR-iPSC reporter line), PR – photoreceptor, cARR3 – cone arrestin (cone PR marker), scale bar 50µm*

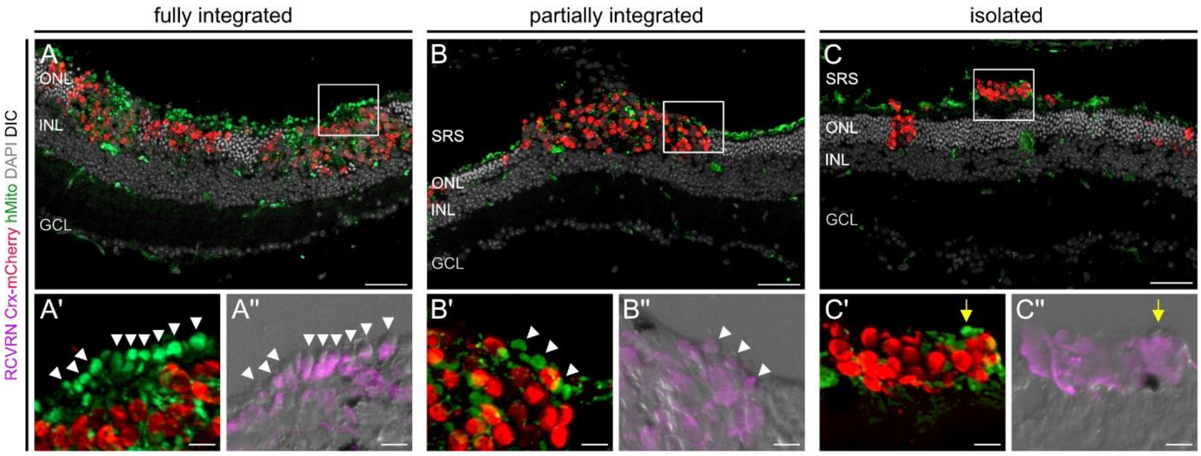


***Suppl. Figure 3: Categorization of graft incorporation***

*Exemplary images of fully (A), partially (B) integrated, and isolated (C) photoreceptor grafts (TP D120; mCherry - red). Inner segments labeled with hMito (human-specific mitochondria marker (green, depicted by white arrowheads) that polarize apically towards the SRS are mainly found in fully integrated transplants and to less propensity in partially integrated grafts. hMito false positive structures (marked by yellow arrow) were excluded for quantification. hMito co-localizes with RCVRN (magenta) but not Crx-mCherry, additionally confirming human PR identity and polarization.*

*TP – Tenpoint protocol, PR – photoreceptor, IS – inner segment, SRS – subretinal space, ONL – outer nuclear layer, INL – inner nuclear layer, GCL – ganglion cell layer, scale bar: 50µm, blow-ups: 10µm*

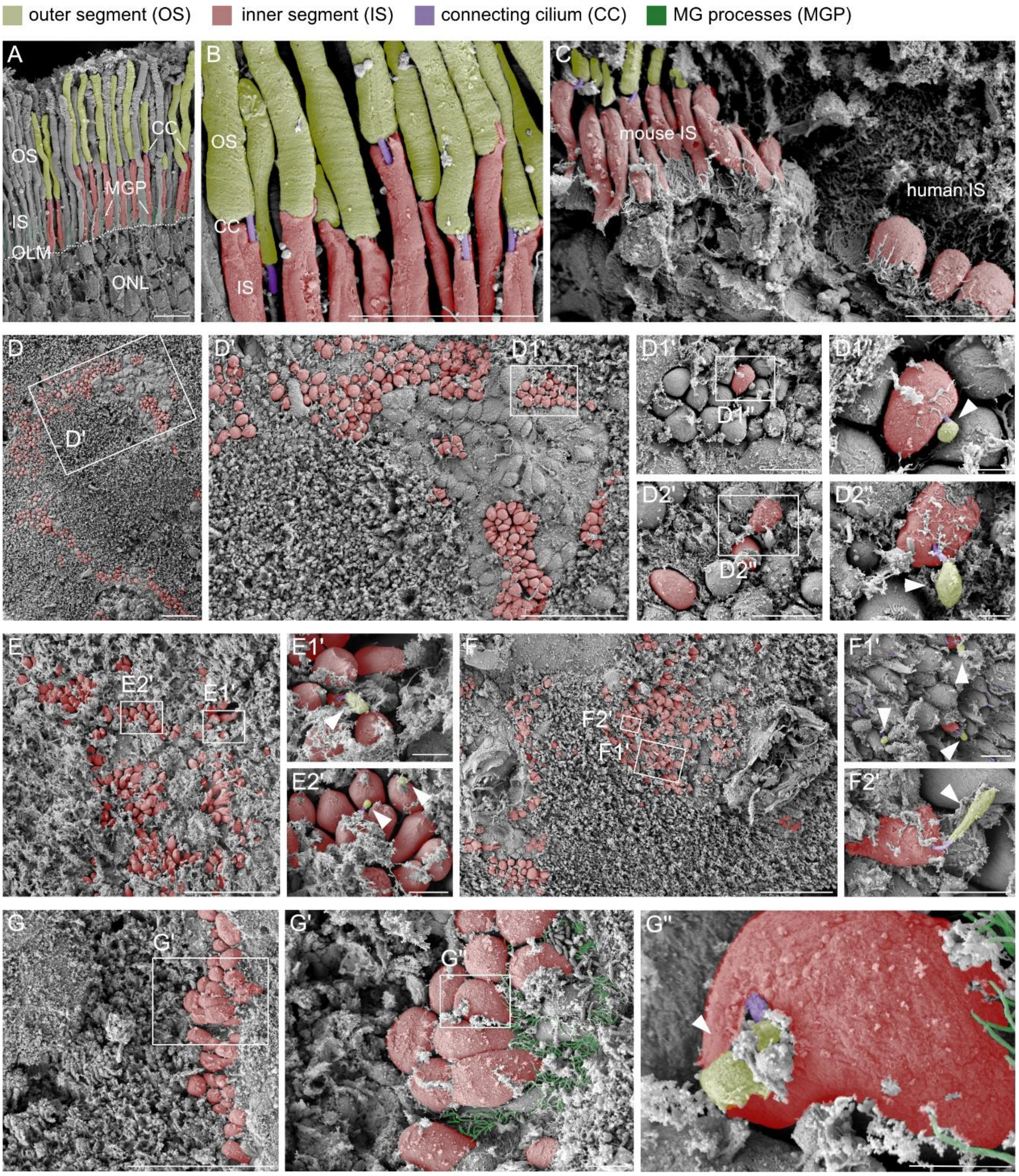


***Suppl. Figure 4: En-face SEM images reveal formation of inner- and outer-segments***

*Exemplary scanning electron microscopic (SEM) images (A-B) of age-matched Cpfl1 mouse retina (36wks, non-transplanted region) illustrate normal retinal structure with intact OLM (dashed line), Müller glia processes (MGP; green) wrapping up PR IS (red), and normal PR ultra-structure including IS (red) and OS (yellow-green) connected by CC (magenta). C) shows the structure of host and donor PR side-by-side (D120, 6M post-tp) (OS, IS, and CC colored as indicated), with mouse IS being long and slender, whereas human IS of transplanted PR are short and stumpy.*

*En face SEM images of the retinal boundary of eyes transplanted with D120 PR (D-D2''), D150 PR (E-E2'), D200 PR (TP; F-F2'), and D200 PR (CTRL; G-G'') illustrate the formation of OS equivalents (yellow-green) that are correctly connected to IS (red) via connecting cilia (magenta), in all experimental paradigms. N (transplantation rounds), n (transplanted eyes) assessed per experimental paradigm: A) D120 (TP) N=1, n=1; B) D150 (TP) N=1, n=1; C) D200 (TP) N=1, n=1; D) D200 (CTRL) N=1, n=1. If not state otherwise, donor source are HROs generated with the Tenpoint protocol.*

*OS – outer segments, IS – inner segments, CC – connecting cilium, MGP – Müller glia processes, D – day of culture, PR – photoreceptors, M – month, wks – weeks, post-tp – post transplantation, scale bars: 50µm (D, D', E, F, G), 10µm (A, B, C, D1', D2'), 5µm (E1', E2', F1', F2', G'), 2µm (D1'', D2'', G'').*

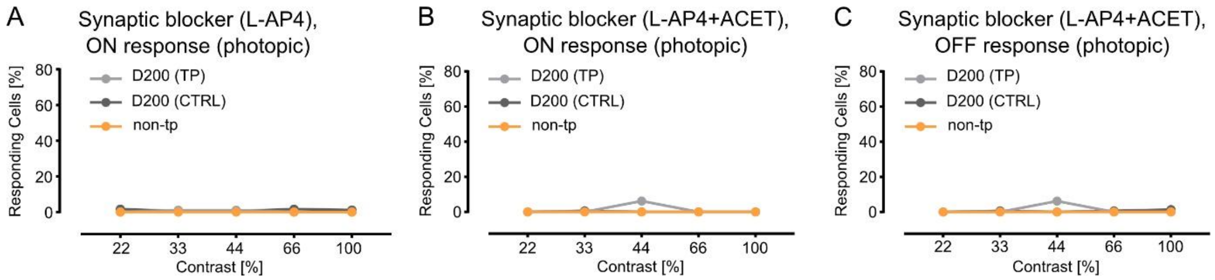


***Suppl. Figure 5: Effective pharmacological inhibition of ON and OFF pathways***

*Incubation with the synaptic inhibitors L-AP4 and ACET results in complete blocking of ON (A,B) and OFF (C) pathways, respectively, thus excluding contribution of intrinsically photosensitive RGCs to measured spiking. N (transplantation rounds), n (transplanted eyes) used per experimental paradigm: A-C) D200 (TP) N=2, n=7; D200 (CTRL) N=2, n=6, non-tp – non-transplanted (age-matched control) n=6.*

*TP – Tenpoint, CTRL – control, non-tp – non-transplanted (age-matched control), L-AP4 – L-2-amino-4-phosphonobutyric acid, ACET – (S)-1-(2-Amino-2-carboxyethyl)-3-(2-carboxy-5-phenylthiophene-3-yl-methyl)-5-methylpyrimidine-2,4-dione*
